# The dual resistance mechanism of CYP325G4 and CYP6AA9 in *Culex pipiens pallens* legs according to transcriptome and proteome analysis

**DOI:** 10.1101/2024.05.13.593821

**Authors:** Yang Xu, Jiajia Du, Kewei Zhang, Jinze Li, Feifei Zou, Xixi Li, Yufen Meng, Yin Chen, Li Tao, Fengming Zhao, Lei Ma, Bo Shen, Dan Zhou, Yan Sun, Guiyun Yan, Changliang Zhu

## Abstract

Mosquitoes within the *Culex pipiens* complex play a crucial role in human disease transmission. Insecticides, especially pyrethroids, are used to control these vectors. Mosquito legs are the main entry point and barrier for insecticides to gain their neuronal targets. However, the resistance mechanism in mosquito legs is unclear. Herein, we employed transcriptomic analyses and isobaric tags for relative and absolute quantitation techniques to investigate the resistance mechanism, focusing on *Cx. pipiens* legs. We discovered 2346 differentially expressed genes (DEGs) between deltamethrin-resistant (DR) and deltamethrin-sensitive (DS) mosquito legs, including 41 cytochrome P450 genes. In the same comparison, we identified 228 differentially expressed proteins (DEPs), including six cytochrome P450 proteins). Combined transcriptome and proteome analysis revealed only two upregulated P450 genes, *CYP325G4* and *CYP6AA9*. The main clusters of DEGs and DEPs were associated with metabolic processes, such as cytochrome P450-mediated metabolism of drugs and xenobiotics. Transcription analysis revealed high *CYP325G4* and *CYP6AA9* expression in the DR strain at 72 hours post-eclosion compared with that in the DS strain, particularly in the legs. Mosquitoes knocked down for *CYP325G4* were more sensitive to deltamethrin than the controls. *CYP325G4* knockdown reduced the expression of several chlorinated hydrocarbon (CHC)-related genes, which altered the cuticle thickness and structure. Conversely, *CYP6AA9* knockdown increased CHC gene expression without altering cuticle thickness and structure. P450 activity analysis demonstrated that CYP325G4 and CYP6AA9 contributed to metabolic resistance in the midgut and legs. This study identified CYP325G4 as a novel mosquito deltamethrin resistance factor, being involved in both metabolic and cuticular resistance mechanisms. The previously identified CYP6AA9 was investigated for its involvement in metabolic resistance and potential cuticular resistance in mosquito legs. These findings enhance our comprehension of resistance mechanisms, identifying P450s as promising targets for the future management of mosquito vector resistance, and laying a theoretical groundwork for mosquito resistance management.

**Author Summary:** *Culex pipiens* mosquitoes are the primary vector of the filamentous nematode, *Wuchereria bancrofti* and also involved in the transmission of other pathogens, such as West Nile virus (WNV), avian malarias, and avian pox virus. Insecticides, particularly pyrethroids, continue to be the primary method to control these significant vectors. Worryingly, resistance to insecticides has become widespread and is rapidly intensifying in *Culex* mosquitoes throughout China, posing a threat to the efficacy of insecticides. Legs are the main sites of contact with ITNs and sprayed insecticides, and the insecticides have to penetrate the leg cuticle to reach their targets.Therefore, the resistance mechanisms in mosquito legs deserve further investigation. Several reports have found a certain amount of P450 in insect legs. Unfortunately, none of the above reports have conducted further functional studies on P450s in the legs. Here, we have identified two P450 enzymes, CYP325G4 and CYP6AA9, through the integrated analysis of transcriptomics and proteomics. CYP325G4 enriched in the cuticle of resistant mosquitoes might possess a dual resistance mechanism involving metabolic resistance and cuticle resistance. CYP6AA9 was slightly different, possibly exerting metabolic resistance as its main function and also being involved in cuticle synthesis. Understanding the dual resistance mechanism of P450s in the metabolism of pyrethroid insecticides will have an important role in optimizing vector control strategies.

## Introduction

Mosquitoes are important insect vectors of diseases such as West Nile Virus, Zika, yellow fever, dengue fever, and, especially, malaria, posing a major threat to global human health [1–5]. Worldwide, there were approximately 56 million cases of dengue and 229 million cases of malaria (with 409,000 deaths) in 2019 [6]. In the last five decades, there has been an approximately 30-fold increase in the incidence of dengue fever worldwide, and a recent study suggested that different species of mosquitoes will continue to spread globally in the coming decades, with a significant increase in the incidence of dengue fever by 2050 [7, 8]. Chemical control using insecticides is an important means to control mosquito-borne diseases [9]. Pyrethroids are the only insecticides recommended by the World Health Organization for use in mosquito nets. However, resistance to pyrethroid insecticides has increased rapidly in mosquitoes, greatly reducing the efficacy of insecticide-treated bed nets (ITNs) [10–12]. The WHO has called for effective action to delay and prevent the development of insecticide resistance [13, 14]. Worryingly, the lack of in-depth understanding of the molecular mechanisms of resistance presents a major challenge to develop strategies to manage resistance to pyrethroids [15].

The mechanisms of insecticide resistance include three main aspects. Firstly, target resistance, which is not sensitive to target sites, is caused by structural modifications or mutations (point mutations) of genes that encode target proteins that interact with insecticides [16, 17]. The second is metabolic resistance, which enhances the metabolic detoxification of insecticides, including the upregulation or enhanced activity of enzymes such as cytochrome P450s (P450s), carboxyl/cholinesterase (CCE), and glutathione-S transferase (GST), which contribute to heterologous detoxification [18]. The third is cuticle resistance, which is caused by increased deposition of chlorinated hydrocarbons (CHCs) in the stratum corneum or proteins that restrict insecticide penetration [19]. P450s comprise a superfamily of heme thioproteins and are an important class of Phase I detoxifying enzymes that can metabolize both endogenous and exogenous compounds [20]. Cytochrome P450 is encoded by the CYP gene, and studies have shown that multiple cytochrome P450s were involved in mosquito insecticide resistance. It was found that the gene *CYP6P9a/b* was highly expressed in transcriptome sequencing of pyrethroid resistant compared to sensitive malaria mosquitoes [21]. Multiple detoxifying enzymes have been found to be overexpressed in imidacloprid-resistant *Aedes* mosquitoes in Egypt, such as *CYP6BB2*, *CYP9M9*, and *CYP6M11* [22]. Eight P450 genes were found to be upregulated in cypermethrin-resistant *Cx. quinquefasciatus* mosquitoes, including *CYP4C52v1* and *CYP6BY3* [23]. These findings not only emphasized the functional importance of P450s in insecticide resistance, but also revealed that overexpression of P450s was a significant cause of mosquito insecticide resistance.

Herein, we focused on the mosquitoes’ legs because the legs are the main sites of contact with ITNs and sprayed insecticides, and the insecticides have to penetrate the leg cuticle to reach their targets. However, whether metabolic resistance also occurs place in mosquito legs in unknown. The proteome of *Anopheles gambiae* legs did contain detoxification enzymes [24], and a few GSTs and P450s were identified in the transcriptome of tick legs [25]. Compared with that in the whole body, four cytochrome P450s were enriched significantly in the leg of *An. coluzzii*, and seven cytochrome P450s were upregulated in the legs of resistant *An. coluzzii* [26]. Recently, a dataset comprising *Drosophila melanogaster* single-cell transcriptomic data was shown to contain transcripts of detoxification enzymes, including CYP450s, in cell types from the legs [27]. Unfortunately, none of the above reports have conducted further functional studies on P450s in the legs.

Therefore, in this study, we aimed to determine the molecular mechanism of mosquito leg insecticide resistance using transcriptomics (RNA sequencing (RNA-seq)) and proteomics (isobaric tags for relative and absolute quantitation (iTRAQ)).

## Results

### Leg transcriptome

Fig 1 shows a flow chart of the experiment. A total of 2346 DEGs were obtained in the comparison of the transcriptomes between the DR and DS mosquito legs (Fig 2A), which were associated with metabolic processes, including drug metabolic-Cytochrome P450 and metabolism of xenobiotics (Fig 2D). These DEGs included 41 p450 genes (Fig 2B)(14 upregulated). Among them, two genes, *CYP325G4* and *CYP6AA9* showed significant overexpression in DR legs compared with that in DS legs (Fig 2C).

**Fig 1.**
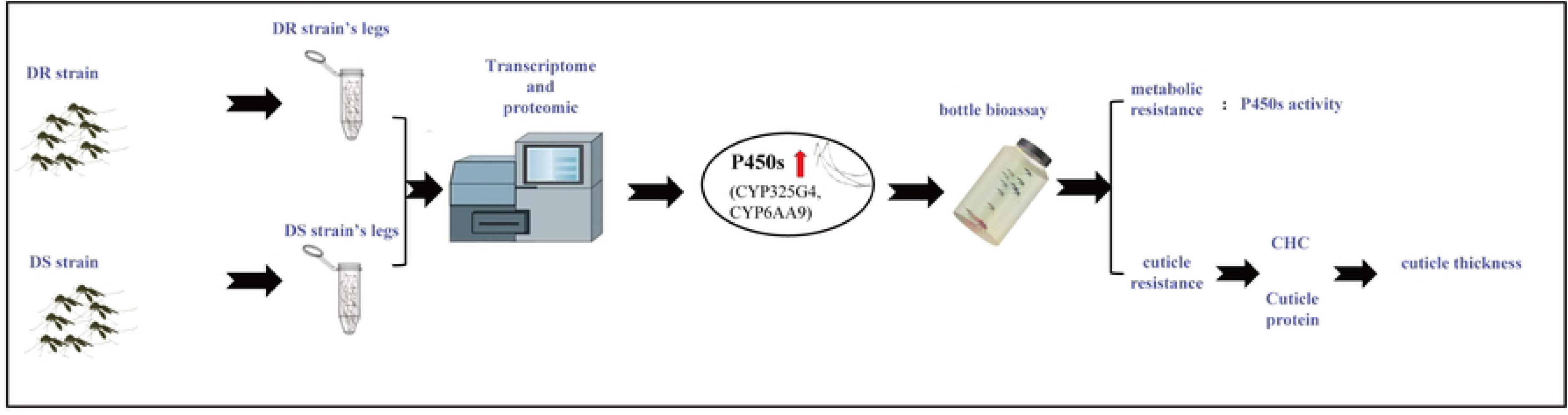
Experimental flowchart of the study. DR, deltamethrin-resistant; DS, deltamethrin-sensitive; CHC, chlorinated hydrocarbon.

**Fig 2.**
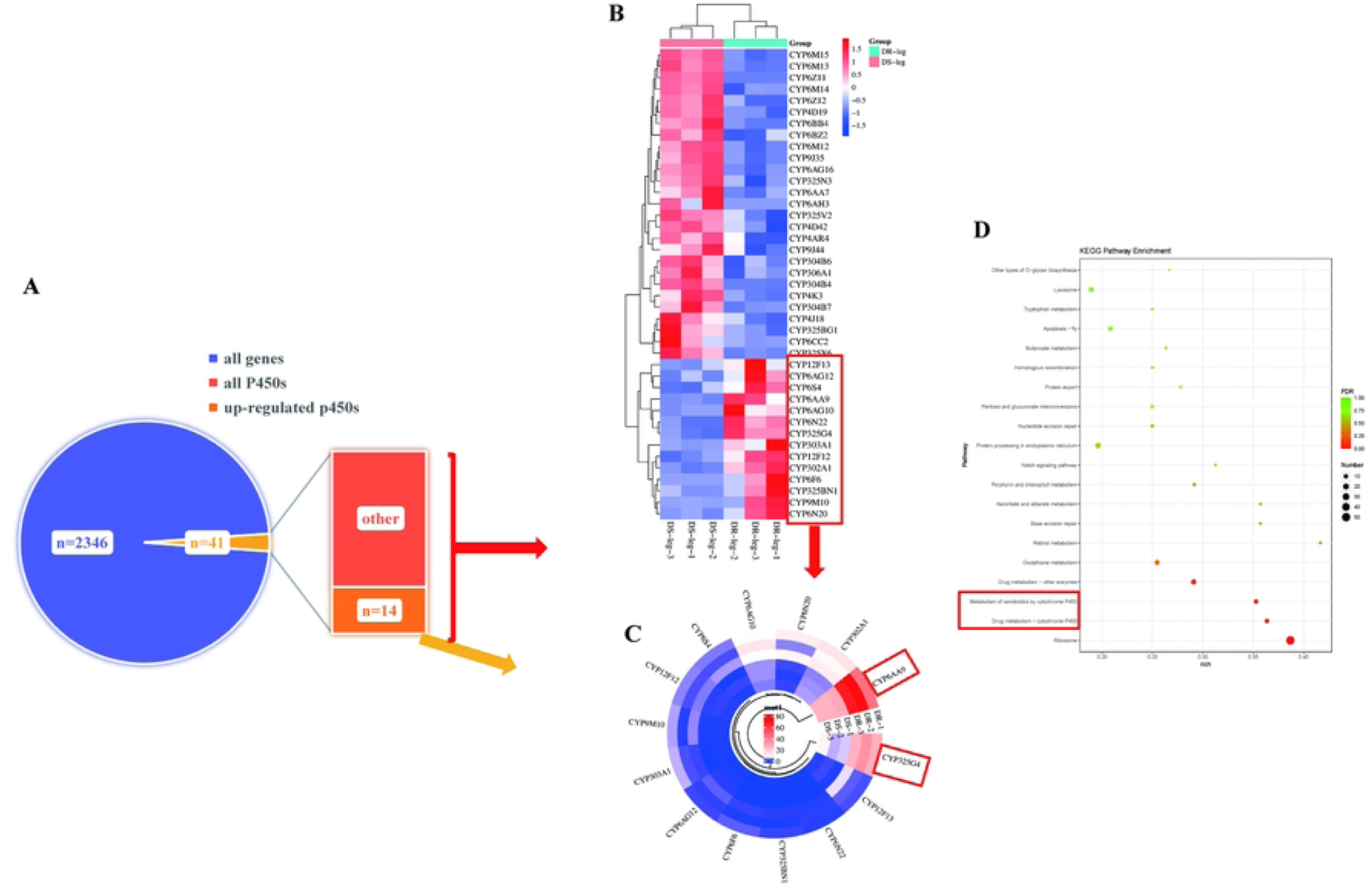
Genes that were up- or downregulated between DR and DS legs in the RNA-seq experiments. **A** Functional classification of 2346. **B** DEGs. Z-scores (normalized expression levels) for the differential expression of cytochrome P450 and upregulated P450s. **C** The top 10 pathways of the DEGs, identified using KEGG enrichment analysis. DR, deltamethrin-resistant; DS, deltamethrin-sensitive; RNA-Seq, RNA sequencing; KEGG, Kyoto Encyclopedia of Genes and Genomes; DEG, differentially expressed gene.

### Leg proteome

A total of 228 differentially expressed proteins DEPs were obtained between the DR and DS leg samples (Fig 3A), which were associated with metabolic processes, including drug metabolic-Cytochrome P450 and metabolism of xenobiotics (Fig 3C). These DEPs included six p450 proteins, two of which were upregulated: CYP325G4 and CYP6AA9 (Fig 3B).

**Fig 3.**
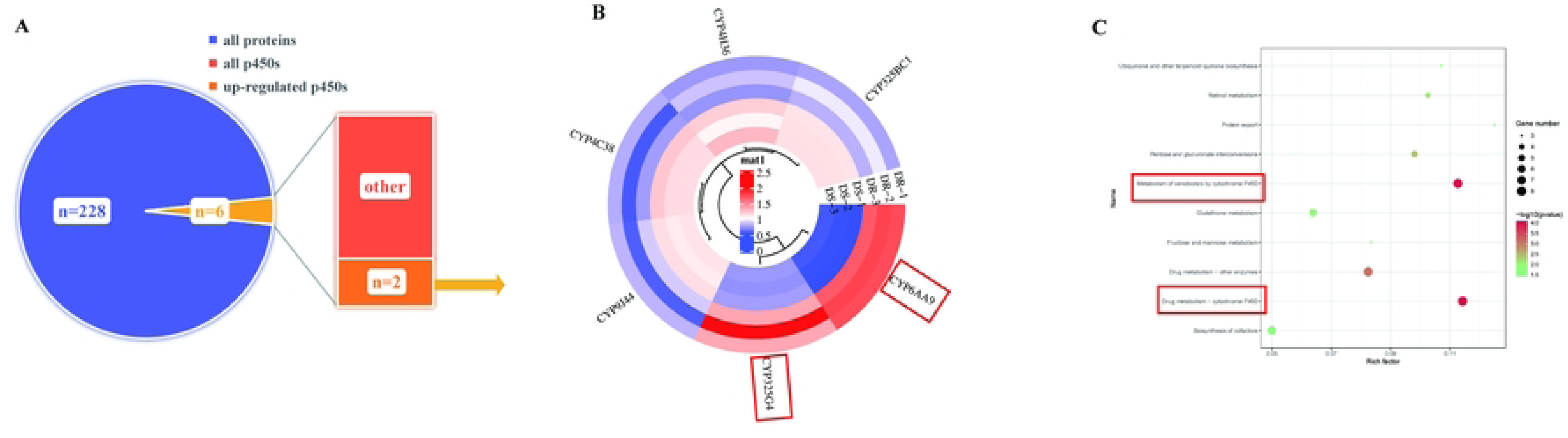
Significantly up- or downregulated proteins between the DR and DS legs. **A** Functional classification of 228 DEPs. Normalized expression levels (z-scores) for **B** differential expression of cytochrome P450 proteins. **C** The top 10 pathways of the DEPs, identified using KEGG enrichment analysis. DR, deltamethrin-resistant; DS, deltamethrin-sensitive; KEGG, Kyoto Encyclopedia of Genes and Genomes; DEP, differentially expressed protein.

### Correlation between transcripts and proteins

We identified 59 proteins that were differentially expressed at both the mRNA and protein levels between the DR legs and DS legs (Fig 4A), which were associated with metabolic processes, including drug metabolic-Cytochrome P450 and metabolism of xenobiotics (Fig 4C). Among them, the only upregulated P450s were CYP325G4 and CYP6AA9 (Fig 4B).

**Fig 4.**
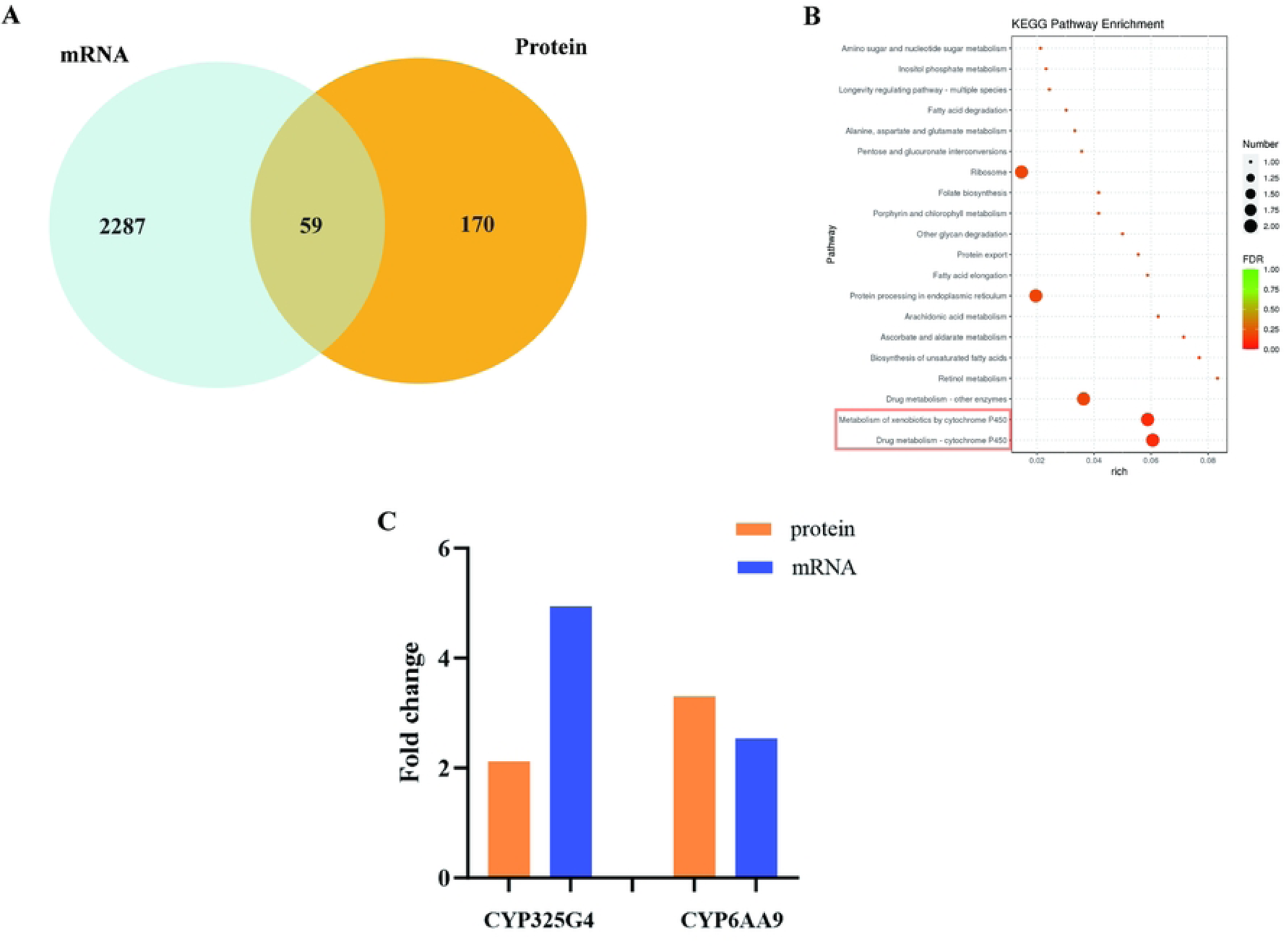
Summary of CYPs that were overexpressed in transcriptomic and proteomic analyses. **A** Venn diagrams depicting the distinctions between different transcripts and proteins. **B** The top 20 pathways of the DEGs and DEPs, identified using KEGG enrichment analysis. **C** Upregulation of two CYP450s at the transcript and protein levels. DEG, differentially expressed gene; DEP, differentially expressed protein; KEGG, Kyoto Encyclopedia of Genes and Genomes.

### CYP325G4 and CYP6AA9 are overexpressed in the DR strain and in mosquito legs

The expression levels of *CYP325G4* and *CYP6AA9* in the DR strain were 3.7-fold (*P* = 0.0079; Fig 5A) and 35.5-fold (*P* = 0.0136)(Fig 5B) higher than those in the DS strain, respectively. The expression levels of *CYP325G4* and *CYP6AA9* in various tissues from female mosquitoes at 72 h PE were examined using qRT-PCR. *CYP325G4* and *CYP6AA9* expression levels were enriched in the legs, and were highly expressed in DR legs compared with that in DS legs (Fig 6A,6B), suggesting that CYP325G4 and CYP6AA9 have important functions in mosquito legs.

**Fig 5.**
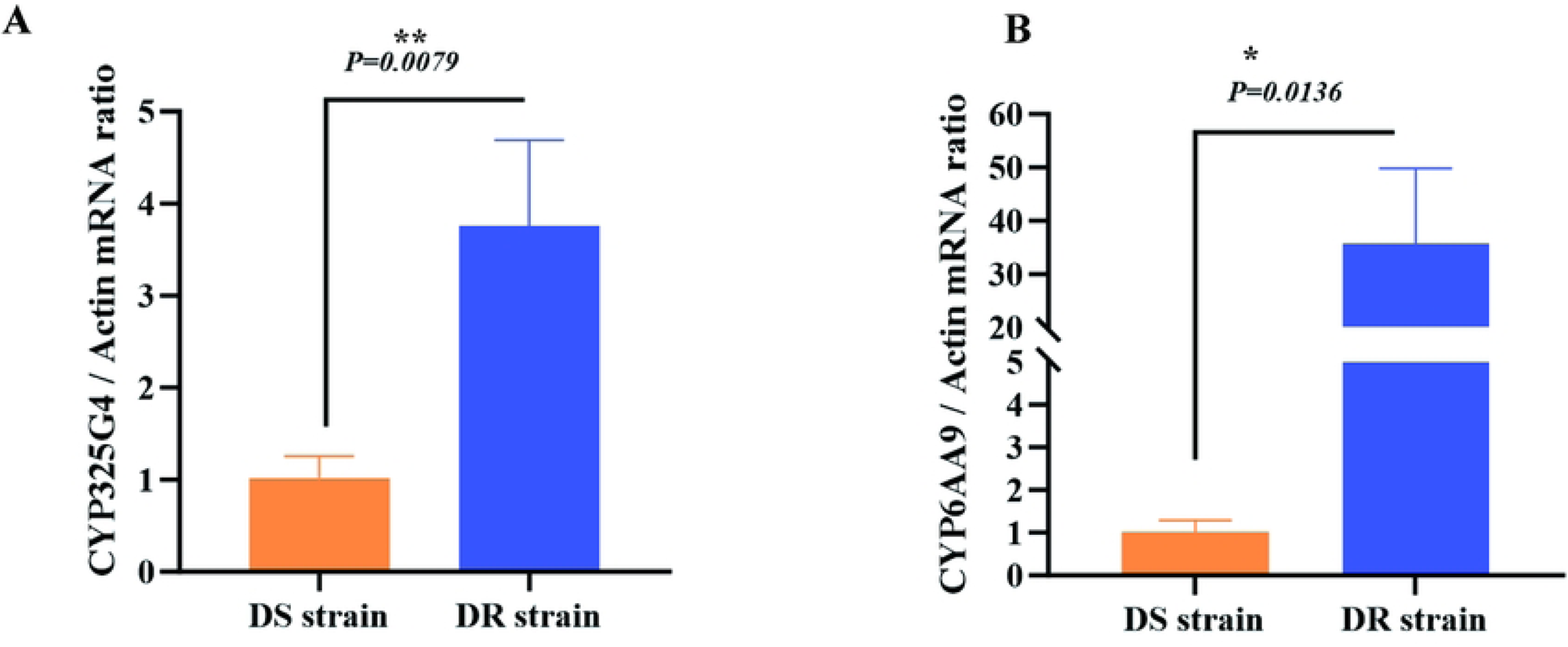
*CYP325G4* and *CYP6AA9* expression in DR and DS mosquitoes. **A** *CYP325G4* expression levels at 72 h PE in the DR and DS strains. **B** *CYP6AA9* expression levels at 72 h PE in the DR and DS strains. The lowest expression value was assigned an arbitrary value of 1, which was then used to calculate the relative expression levels. The results appear as the mean ± standard deviation (SD) (n = 3 biological replicates). \**P* ≤ 0.05; \*\**P* ≤ 0.01. PE, post-eclosion; DR, deltamethrin-resistant; DS, deltamethrin-sensitive.

**Fig 6.**
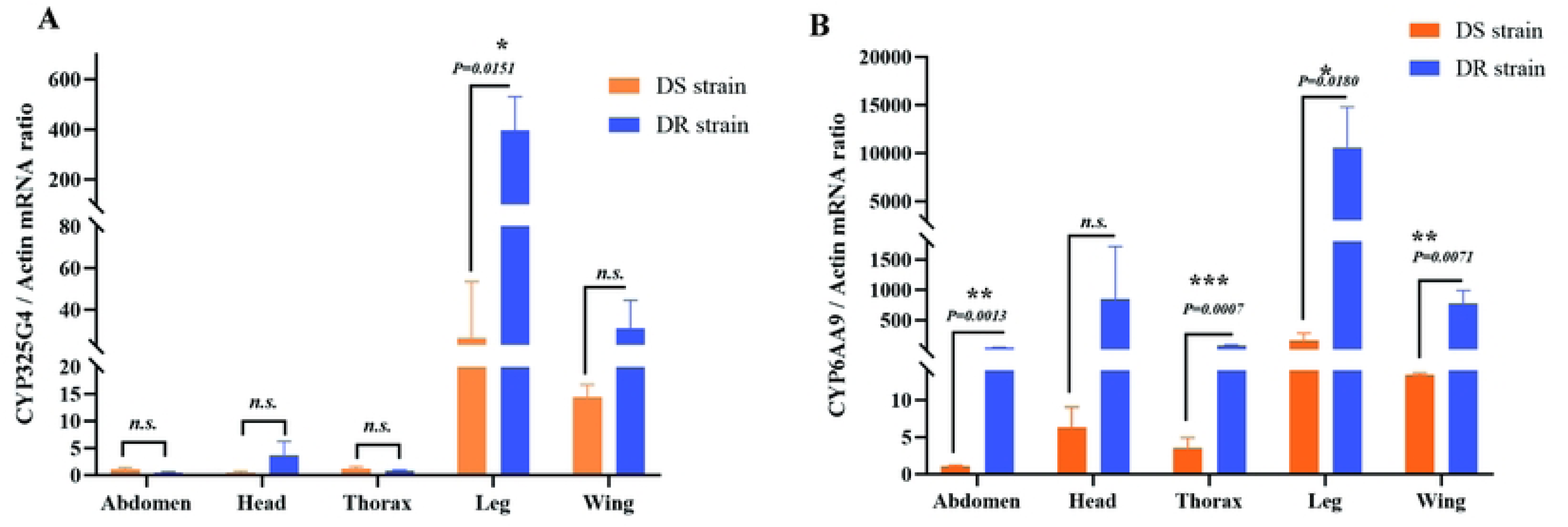
*CYP325G4* and *CYP6AA9* expression profiles in various mosquito tissues. **A** Constitutive *CYP325G4* expression in the DR and DS strains. **B** Constitutive *CYP6AA9* expression in the DR and DS strains. The mRNA levels in the head, thorax, abdomen, legs, and wings of the DS and DR mosquitoes were determined. The lowest expression value was assigned an arbitrary value of 1, which was then used to calculate the relative expression levels. The results appear as the mean ± standard deviation (SD) (n = 3 biological replicates). \**P* ≤ 0.05; \*\**P* ≤ 0.01, \*\*\*\**P* ≤ 0.0001; ns, not significant; DR, deltamethrin-resistant; DS, deltamethrin-sensitive.

### The role of CYP325G4 and CYP6AA9 in deltamethrin resistant mosquitoes

Subsequently, RNAi was used to knockdown the expression of *CYP325G4* and *CYP6AA9*, separately, and functional analysis was performed. The interference efficiency of *CYP325G4* was 65% (*P* < 0.0001; Fig 7A) for the whole body and 54% (*P* = 0.0017; Fig 7B) for the legs. The interference efficiency of *CYP6AA9* was 37% (*P* = 0.0059; Fig 7C) for the whole body and 36.6% (*P* = 0.0087; Fig 7D) for the legs. The CDC bottle biology assay of the siCYP325G4 group revealed that exposure to deltamethrin enhanced mosquito mortality rates from 15 to 120 minutes compared with that of the control group (*P* < 0.05). At 120 minutes of exposure to deltamethrin, the siCYP325G4 group had a 48.1% higher mortality rate than the control group (*P* < 0.05; Fig 8). Thus, silencing of *CYP325G4* enhanced mosquito susceptibility to deltamethrin, indicating that CYP325G4 was implicated in mosquito deltamethrin resistance. As reported previously, *CYP6AA9* knockdown enhanced the sensitivity of adult female mosquitoes to deltamethrin [28].

**Fig 7.**
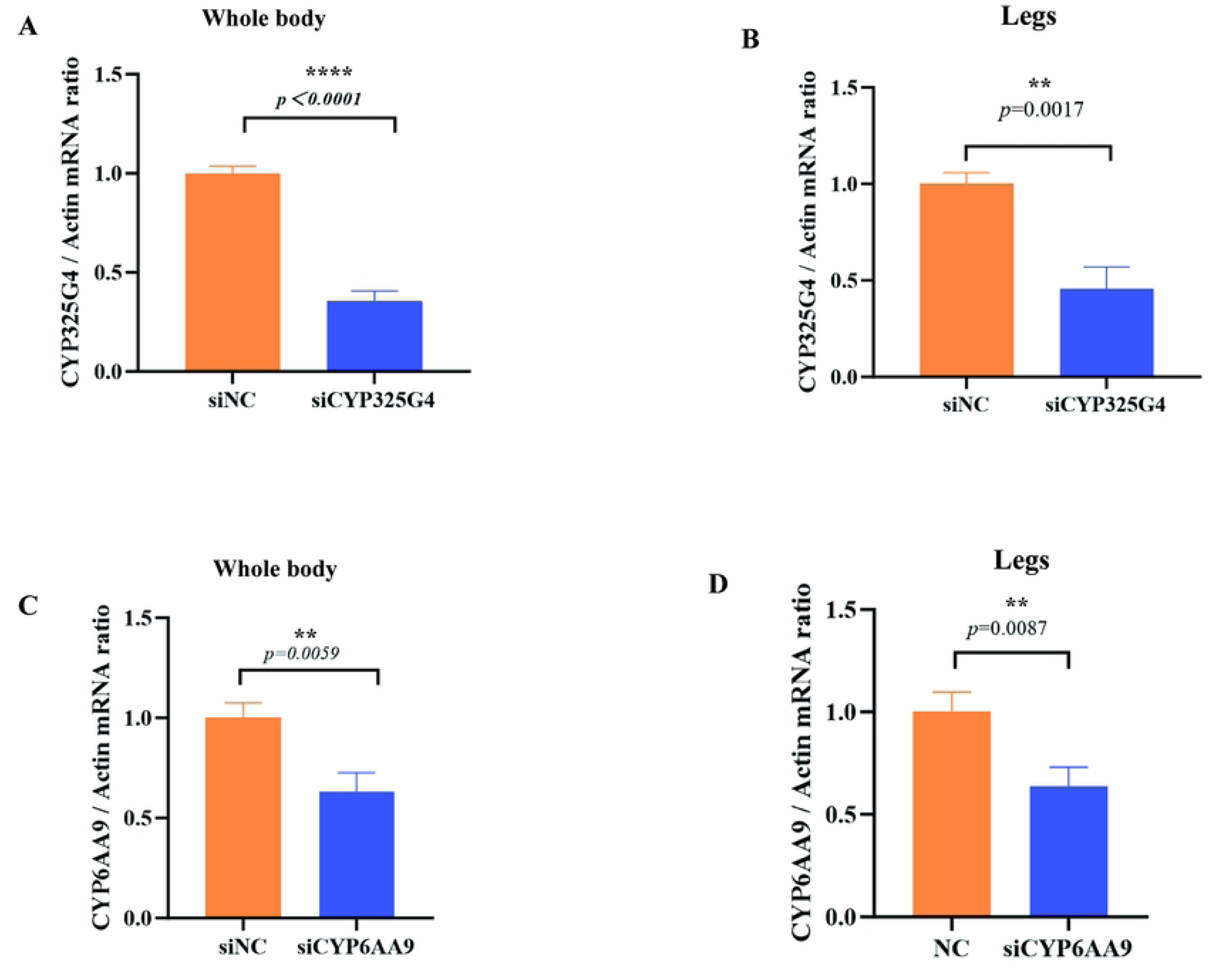
Relative expression levels of *CYP325G4* and *CYP6AA9* after RNAi. *CYP325G4* expression in whole mosquito bodies **A** and legs **B** after CYP325G4 silencing. *CYP6AA9* expression in whole mosquito bodies **C** and legs **D** after *CYP6AA9* silencing. The results appear as the mean ± standard deviation (SD) (n = 3 biological replicates). \**P* ≤ 0.05; \*\**P* ≤ 0.01. RNAi, RNA interference; siNC, negative control small interfering RNA; siCYP325G4, small interfering RNA targeting *CYP325G4*; siCYP6AA9, small interfering RNA targeting *CYP6AA9*.

**Fig 8.**
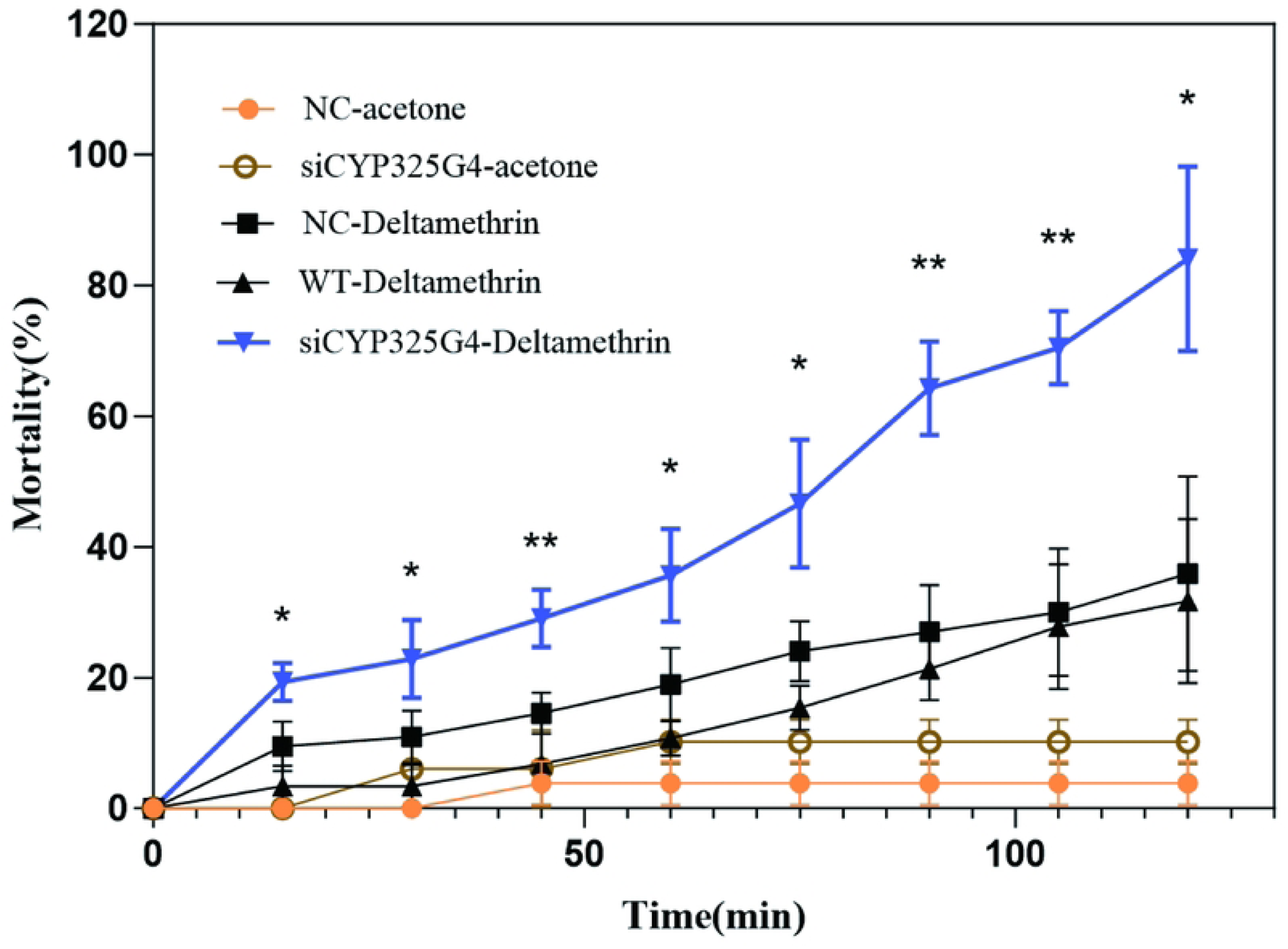
CDC bottle bioassay after *CYP325G4* knockdown in the DR strain. CDC bottle assays (0.1 mg/ml) to determine the level of insecticide resistance following *CYP325G4* silencing. The results appear as the mean ± standard deviation (SD) (n = 3 biological replicates). The p-values were determined relative to the results in the NC group. The results for the DEPC and NC injected groups were not significantly different. \**P* ≤ 0.05; \*\**P* < 0.01. CDC, Center for Disease Control; DR, deltamethrin-resistant; siNC, negative control small interfering RNA; WT, wild-type; siCYP325G4, small interfering RNA targeting *CYP325G4*; DEPC, diethyl pyrocarbonate.

### Silencing *CYP325G4* and *CYP6AA9* affected the expression of genes related to hydrocarbon synthesis

We further explored the resistance role exerted by CYP325G4 and CYP6AA9 in the leg. The formation of the mosquito cuticle is closely connected with the synthesis of chlorinated hydrocarbons (CHCs). First, we investigated whether *CYP325G4* and *CYP6AA9* silencing influenced the expression of six key CHC synthesis genes. According to qRT-PCR analysis, knockdown of *CYP325G4* decreased the mRNA expression level of acetyl-coenzyme A carboxylase (ACC), fatty acid synthase S-acetyltransferase (FAS), elongase (CPIJ003715), very-long-chain fatty acid-coenzyme A ligase (FACVL), and CYP303A1 in the whole body (Fig 9A). In the legs, knockdown of *CYP325G4* efficiently decreased the mRNA expression level of elongase (CPIJ015710), elongase (CPIJ003715), FACVL, and CYP303A1(Fig 9B). In the whole body, silencing of *CYP6AA9* increased the mRNA expression of ACC and FAS (Fig 9C), and increased the mRNA expression of ACC, FAS, elongase (CPIJ003715), and elongase (CPIJ015710) in the legs (Fig 9D). We proposed that CYP325G4 and CYP6AA9 affected the expression of CHC biosynthesis genes via different regulatory pathways, and thus their potential functions might be different.

**Fig 9.**
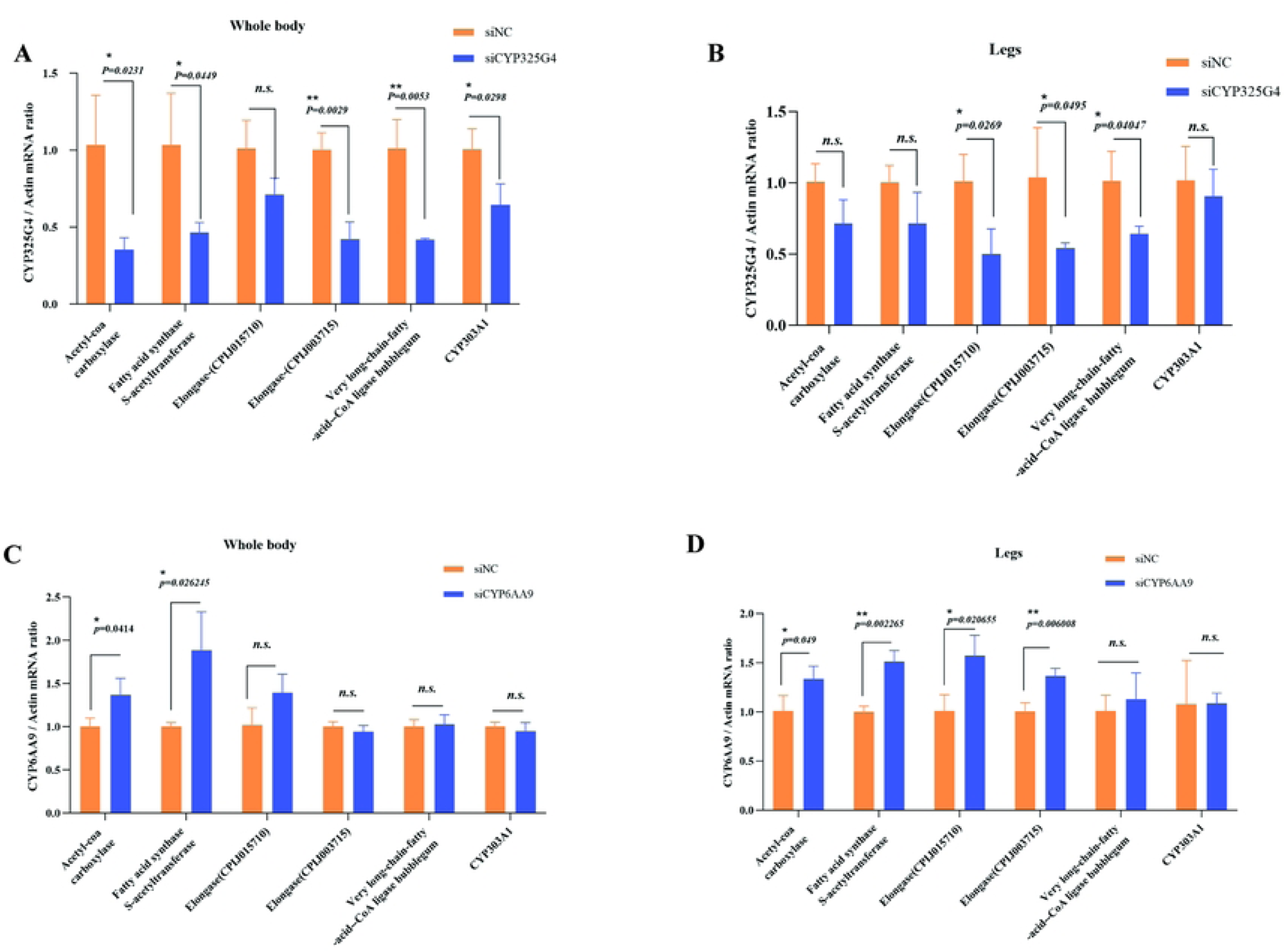
Expression levels of CHC-related genes after microinjection of siCYP325G4 and siCYP6AA9. **A** Expression levels of CHC-related genes in whole mosquitoes after microinjection of siCYP325G4; **B** Expression levels of CHC-related genes in the legs after microinjection of siCYP325G4. **C** Expression levels of CHC-related genes in whole mosquitoes after microinjection of siCYP6AA9; **D** Expression levels of CHC-related genes in the legs after microinjection of siCYP6AA9. \**P* ≤ 0.05; \*\**P* ≤ 0.01; ns, not significant. CHC, chlorinated hydrocarbon; siNC, negative control small interfering RNA; siCYP325G4, small interfering RNA targeting *CYP325G4*; siCYP6AA9, small interfering RNA targeting *CYP6AA9*.

### SEM analysis of cuticle thickness

Subsequently, we analyzed mosquito legs using SEM, which revealed uneven cuticle thickness in the RNAi group. The siCYP325G4 group had a lower average cuticle thickness (1.349 ± 0.29µm) than the siNC group (1.676 ± 0.25µm; *P* = 0.0185)(Fig 10A,B). The result implied that CYP325G4 might cause cuticle resistance by influencing the CHC content, which would lead to cuticle changes in mosquito legs. Compared with that in the control group, there was no change in the cuticle structure and thickness in the siCYP6AA9 group (Fig 10C,D; Table 1).

**Fig 10.**
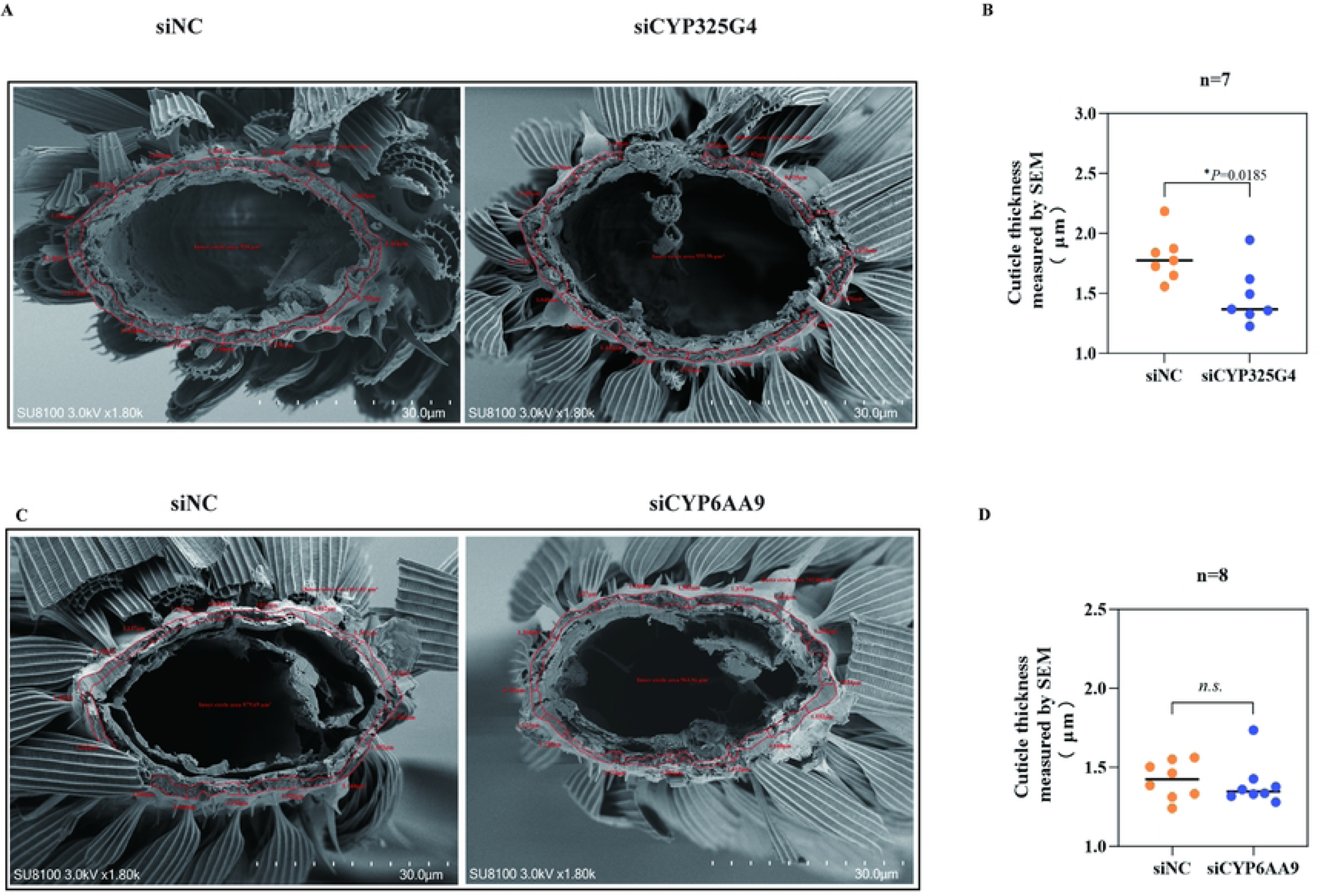
Leg cuticle thickness in mosquitoes treated with siCYP325G4 and siCYP6AA9. **A** SEM images of the front view of a sectioned leg from the siNC group and siCYP325G4 group. **B** Column bar graph (vertical) of the whole cuticle thickness after microinjection of siNC and siCYP325G4. **C** SEM images of the front view from a sectioned leg of the siNC group and the siCYP6AA9 group. **D** Column bar graph (vertical) of the whole cuticle thickness after microinjection of siNC and siCYP6AA9. The mean cuticle thickness was calculated using 16 points of measurement in each individual. Results appear as the mean ± SD. n = the number of measurements performed for each batch of 7 or 8 mosquitoes. \**P* ≤ 0.05; ns, not significant. SEM, scanning electron microscope; siNC, negative control small interfering RNA; siCYP325G4, small interfering RNA targeting *CYP325G4*; siCYP6AA9, small interfering RNA targeting *CYP6AA9*.

**Table 1.** Average cuticle thickness of each group.

| Group | siNC | siCYP325G4 | siNC | siCYP6AA9 |
| --- | --- | --- | --- | --- |
| Cuticle thickness |  |  |  |  |
| Cuticle thickness by SEM ($\mu\text{m}$) | $1.676 \pm 0.25$ | $1.349 \pm 0.29$ | $1.325 \pm 0.26$ | $1.316 \pm 0.24$ |

### CYP325G4 and CYP6AA9 are involved in mosquito metabolic resistance

To determine whether CYP325G4 and CYP6AA9 are involved in metabolic resistance, P450 enzyme activity was measured in samples from gene-silenced mosquitoes (Fig 11A). The enzyme activities in the siCYP325G4 and siCYP6AA9 groups in the whole body were reduced by 11% (*P* = 0.001) and 10% (*P* = 0.004) relative to that in the control group, respectively (Fig 11B,F), implying that the P450 enzyme activities of CYP325G4 and CYP6AA9 are involved in mosquito metabolic resistance. Additionally, we observed that the P450 enzyme activity of midguts of the siCYP325G4 and siCYP6AA9 group mosquitoes decreased by 81% (*P* < 0.0001) and 19% (*P* = 0.001) respectively (Fig 11C,G). By contrast, the activities in the legs were reduced by 19% (*P* = 0.0382) and 15.7% (*P* = 0.046) respectively (Fig 11D,H) in the siCYP325G4 and siCYP6AA9 groups compared with those in the control group. These results suggested that CYP325G4 and CYP6AA9 exert metabolic resistance, not only in the mosquito midgut, but also in the mosquito leg, which is the initial site of mosquito exposure to insecticides.

**Fig 11.**
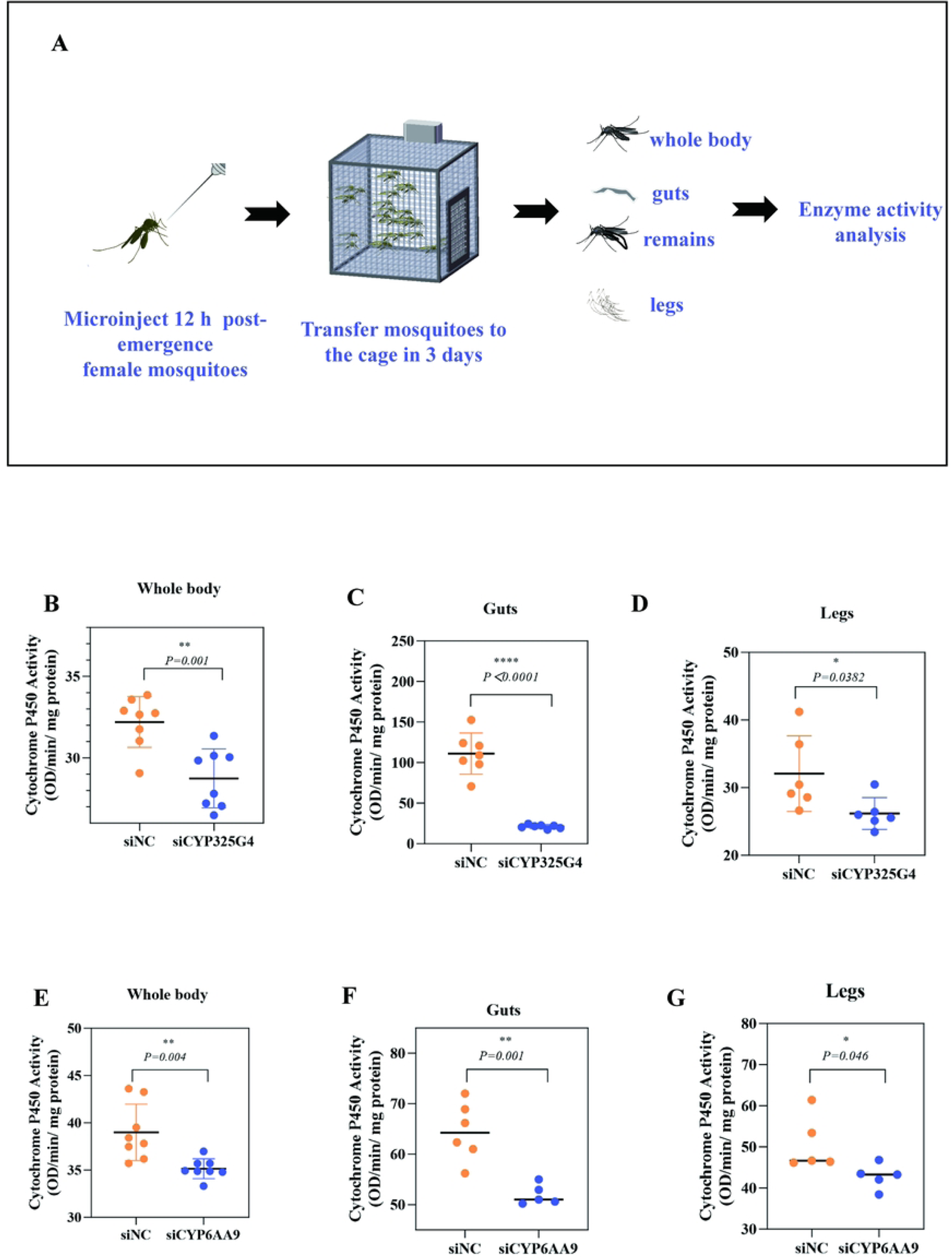
Measurement of cytochrome P450 enzyme activity after microinjection of siCYP325G4 and siCYP6AA9. **A** Schematic diagram of the method used to assess P450 activity. **B** P450 enzyme activities in the whole body after microinjection of siCYP325G4. **C** P450 enzyme activities in the gut after microinjection of siCYP325G4. **D** P450 enzyme activities in the legs after microinjection of siCYP325G4; **E** P450 enzyme activities in the whole body after microinjection of siCYP6AA9; **F** P450 enzyme activities in the guts after microinjection of siCYP6AA9; **G** P450 enzyme activities in the legs after microinjection of siCYP6AA9; \**P* ≤ 0.05; \*\**P* < 0.01;\*\*\*\**P* < 0.0001. siNC, negative control small interfering RNA; siCYP325G4, small interfering RNA targeting *CYP325G4*; siCYP6AA9, small interfering RNA targeting *CYP6AA9*.

## Discussion

### The legs are the primary and important body part related to mosquito resistance

When mosquitoes land on an insecticide-treated net or an insecticide-sprayed surface, their legs are a key point of contact with the insecticide. Therefore, the insecticide must first penetrate the cuticle of the legs to reach the neurons. The cuticle forms the outermost part of the mosquito leg, which exerts various functions, such as movement, mechanical support, perception of the environment, chemical communication, and preventing dryness [29]. The cuticle in the leg also acts as the primary barrier that protect insects from insecticide ingress [29]. In 2019, *Balabanidou* et al. reported that CYP450s were present in the leg-specific proteome [24]. In 2021, the same group reported that four P450s were enriched in the legs of *Anophele*s mosquitoes, and transcriptome analysis identified seven upregulated P450s in pyrethroid-resistant mosquito legs [26]. Several transcripts encoding detoxifying enzymes, such as cytochrome P450, glutathione-S transferase, and UDP glucuronic acid transferase, were identified in recent single-cell transcriptome profiles of *Drosophila melanogaster* [27]. A small number of P450s was also found in the transcriptome of tick legs [25]. Thus, the importance of insect legs in resistance warranted further in-depth study. Indeed, new resistance mechanisms in the legs have been discovered, not only the cuticle resistance caused by cuticle proteins, which leads to cuticle thickening, but also the involvement of chemoreceptors and ABC transporters in the legs of mosquitoes in resistance. Recently, it was found that many P450 gathered in the legs; however, their role was unknown [26, 30]. Our group screened differentially expressed genes and proteins of DR and DS mosquito legs through transcriptome and proteome analysis. We found that multiple P450s were differentially expressed. According to the KEGG enrichment analysis, the drug metabolizing enzyme-cytochrome P450 pathway was enriched in resistant mosquito legs. Nevertheless, it was unclear whether P450s exhibit metabolic resistance in mosquito legs and their exact role was unknown.

### First detection of CYP325G4 and CYP6AA9 expression in mosquito legs

The increased expression of P450s in insect bodies has been related to improved insecticide detoxification levels, evolutionary selection of insecticides, and the ability of insects to adapt to environmental changes [31, 32]. To date, the CYP6 and CYP9 families have been widely reported. Previously, we reported the involvement of CYP6AA9 in mosquito pyrethroid resistance [28]. However, its function remains unexplored. In addition, our group also reported that the *CYP325A3* gene of *An. gambiae* and the *CYP307B1* and *CYP314A1* genes of *Cx. pipiens pallens* are associated with resistance to pyrethroids [33, 34]. *Liu* et al. found that cypermethrin had an inducing effect on the expression of multiple P450s in mosquitoes, with the expression level of the *CYP325G4* gene increasing by more than twice compared with that in response to acetone treatment [32]; however, there was a lack of further mechanistic research on CYP325G4. Herein, we found that cytochrome P450s CYP325G4 and CYP6AA9 were upregulated in both the transcriptome and proteome of the legs of resistant mosquitoes and were both overexpressed in the whole body of resistant mosquitoes.

### CYP325 and CYP6AA9 are not only enriched in legs, but also highly expressed in resistant legs

In recent years, transcriptomic or immuno-localization analysis in local tissues of insecticide-resistant insects (e.g., mosquitoes, *Drosophila*, and *Bemisia tabaci*) has shown that P450s are expressed in many organs and tissues besides the common midgut This tissue specificity of P450s in, for example, the cuticle, oocytes, brain, Malpighian tubules, and Haller’s organ, suggested that the tissue specificity of P450s might be associated with insecticide resistance [35, 36]. Our transcriptome analysis revealed that high expression of multiple P450s in the cuticle of DR *Cx. pipiens pallens*, involving members of the CYP6 family (e.g., AA9, BB3, BB4, M13, and N23), CYP325 family (e.g., BG1, B4, K3, N3, and V2), and CYP4 family (e.g., J13, C38, and D43). Combined transcriptome and proteome analysis showed that CYP325G4 and CYP6AA9 were highly expressed at both the mRNA and protein levels. Analysis using qRT-PCR revealed that *CYP325G4* and *CYP6AA9* were not only enriched in legs, but also highly expressed in the legs of resistant mosquitoes, suggesting that CYP325G4 and CYP6AA9 might play an important role in insecticide resistance in the legs.

### Cuticle resistance involving CYP325G4 and CYP6AA9

Recent studies have found that CYP450s not only play a metabolic detoxification role in insects, but also have an important function in cuticle formation. CYP4G16 was found to confer mosquito insecticide resistance through the formation of catalytic CHCs in a multi-insecticide resistant strain of *An. gambiae* [37]. In *Locusta migratoria*, researchers found that follicle-cell-specific CYP450 *Lm*CYP4G102 was inhibited, resulting in a dramatic decrease in the total alkane content and a significant increase in cuticular fragility [38]. In *Blattella Germanica*, CYP4G19 is highly expressed in pyrethroid-resistant strains and is involved in the production of CHC, thereby promoting the insect’s ability to resist cuticular penetration [38]. Most studies on P450 genes involved in insect cuticle formation have been related to the CYP4G family. Interestingly, some recent studies have suggested that the CYP3 family might also be involved in insect CHC formation. One of the functions of the *LmCYP303A1* gene in *Locusta migratoria* was to regulate CHC biosynthesis, playing a key role in preventing water loss and insecticide penetration [39]. Research on mosquito CYP genes has mostly focused on metabolic detoxification, with less research on their involvement in cuticle synthesis, especially in the CYP3 family. CHC biosynthesis involves many enzymes [40–42]. In this study, we selected six genes involved in CHC synthesis, including those encoding ACC, FAS, two elongation enzymes, FACVL, and CYP303A1. mRNA level analysis showed that all six genes were decreased to varying degrees after RNAi of *CYP325G4*. However, knockdown of *CYP6AA9* efficiently promoted the expression of the several CHC genes. Thus, CYP325G4 and CYP6AA9 showed opposite regulatory trends. We speculated that CYP325G4 and CYP6AA9 affect CHC biosynthesis-related gene expression via different regulatory pathways and their potential functions might be different.

Finally, we observed the cuticle structure of mosquito legs using SEM. The control group had a cuticle with uniform thickness and a dense structure, while the siCYP325G4 group showed uneven thickness and thinning of the tarsal cuticle, which might benefit insecticide penetration. These results suggested that cuticle-enriched CYP325G4 in resistant mosquitoes might participate in cuticle resistance by changing the structure or composition of the cuticle via CHCs. However, compared with the control group, there was no change in the cuticle structure and thickness of the siCYP6AA9 group. This result was very complex and required further research.

### Metabolic resistance of CYP325G4 and CYP6AA9 in the legs and midgut

Multiple studies have shown that an increase in P450 expression levels leads to an increase in total P450 levels, as well as an increase in P450 enzyme activity, resulting in insecticide resistance [23, 43, 44]. The CYP450 enzyme activity of resistant *An. stephensi* in different regions was 2.23 and 2.54-fold higher than that of sensitive strains [45]. The activity of monooxygenases in pyrethroid resistant *An. stephensi* was 1.88-fold higher than that of sensitive strains [46]. In this study, we found that RNAi of *CYP325G4* enhanced mosquito susceptibility to deltamethrin, suggesting that CYP325G4 is associated with insecticide resistance. CYP6AA9 has already been reported to be related to resistance [28]. The activities of P450s decreased after *CYP325G4* and *CYP6AA9* silencing. This suggested that CYP325G4 and CYP6AA9 are involved in metabolic resistance.

We found that the RNAi groups had the most significant decreases in P450 enzyme activities in the midgut compared with those of the siNC group. Notably, compared with that in the legs of the siNC group, the legs of the RNAi groups showed a significant decrease in P450 enzyme activity, suggesting that CYP325G4 and CYP6AA9 not only exerted metabolic resistance in the common metabolic site of the mosquito midgut, but also in the initial site of mosquito contact with insecticides, i.e., the legs. We speculated that mosquito legs are involved in the immediate degradation and metabolism of insecticides as the first line of defense. Metabolizing insecticides before they enter the mosquito body and exert their toxic effects, would ameliorate at least some of the toxic effects of insecticides, and even protect the peripheral nerves of the legs from pyrethroid toxicity. This study is the first report of metabolic insecticide resistance mediated by CYP450s not only in the guts but also in the legs.

## Conclusions

As the first line of defense, the thickened cuticle of mosquito legs could reduce the penetration of certain insecticides. Insecticides that have already penetrated into the leg cuticle could be partially degraded by leg-resident CYP325G4 and CYP6AA9. When the insecticide reaches the midgut of mosquitoes, it will be degraded by P450 metabolic enzymes, including CYP325G4 and CYP6AA9, greatly reducing the toxicity of the insecticide to mosquitoes. Consequently, CYP325G4 enriched in the cuticle of resistant mosquitoes might have a dual resistance mechanism of metabolic resistance and cuticle resistance. CYP6AA9 was slightly different, possibly exerting metabolic resistance as its main function and also being involved in cuticle synthesis. Understanding the dual resistance mechanism of P450s in the metabolism of pyrethroid insecticides will have an important role in optimizing vector control strategies.

## Materials and methods

### Strains of mosquito

*Cx. pipiens pallens* mosquitoes (as the deltamethrin-sensitive (DS) strain) were collected from Tangkou (Shandong province, China). This DS strain had a lethal concentration 50 (LC50) for deltamethrin of 0.03 mg/L. Forty generations of selection in DS larvae were used to derive the DR strain (LC50 = 5 mg/L), as reported previously [47].

### Leg dissection

We dissected the whole legs (including the tarsus, tibia, femur, trochanter, and coxa) from non-blood fed 3-day-old female mosquitoes. Three biological replicates (n= 100 legs in each replicate) were used for each strain and/or condition.

### RNA sequencing

A Directzol RNA MiniPrep Plus kit (Zymo Research Corp, Irvine, CA, USA) was used to extract total RNA (100 mosquito legs per tube, three replicates) according to the supplier’s recommendations, including treatment with DNAse. A NanoDrop spectrophotometer (Thermo Fisher Scientific, Waltham, MA, USA) was used to assess the RNA purity and concentration, and a Bioanalyzer (Agilent, Santa Clara, CA, USA) was used to confirm the RNA integrity. Three replicates of each genotype were analyzed. PGM sequencing carried out the RNA-seq protocol at the UTHSC Molecular Resource Center (MRC, Memphis, TN, USA). The RNA samples were converted into Ion Torrent sequencing libraries followed by processing on 318v2 chips utilizing an Ion Torrent Proton sequencer. Chip-to-chip variability was minimized by sequencing all replicates and groups simultaneously.

### Analysis of RNA-seq data

We chosed the sequence of *Cx. Quinquefasciatus* (https://www.vectorbase.org/organisms/culex-quinquefasciatus) as a reference genome (assembly version: CpipJ1) and a reference gene (geneset version: CpipJ1.3). since there is high similarity between *Cx. pipiens pallens* and *Cx. Quinquefasciatus* [48]. A local Slipstream application running a GALAXY installation was used to align and analyze the RNA-seq data. FASTQC was used to check the quality of the sequencing data using the FASTQ files obtained directly from the sequencer. Nucleotides with a phred score < Q20 were trimmed off the reads. RNA STAR was then employed to align the trimmed FASTQ files to the reference library. Subsequently, read count data for each gene in the reference file were extracted from the generated SAM files. The Transcripts per Million (TPM) method was used to normalize the read counts for the entire experiment. DeSeq2 was then employed to analyze the resulting data. We excluded those genes that did not show a fold change of at least a 1.5 and a p-value > 0.05. The trimmed gene list was then subjected to Benjamini and Hochberg false discovery rate (FDR) correction, retaining genes with an FDR < 0.05. The resultant list of significantly differentially expressed genes (DEGs) was imported into R for visualization. The heatmap.2 function in the gplots R package was used to generate heatmaps [26]. KEGG Automatic Annotation Server (KAAS), (http://www.Jp/kegg/kaas/) was used to annotate sequences of terms and identify the paths involved.

### Quantitative proteomics

To extract the proteins from mosquito legs (100 legs per tube, 3 replicates), the legs of each genotype were homogenized in Radioimmunoprecipitation assay buffer containing a complete Protease Inhibitor Cocktail (Roche, Basel, Switzerland). A Filter Aided Sample Preparation (FASP) column was used to desalt and immobilize the lysates, followed by digestion using trypsin and Tandem Mass Tag (TMT) labeling, as detailed previously [49]. All groups were labeled simultaneously to ensure consistent labeling. Liquid Chromatography with tandem mass spectrometry analysis of the digested labeled peptides was then carried out employing an EASY-nLC 1000 UPLC system (Thermo Fisher Scientific) incorporating a 50 cm EASY-Spray Column coupled with an Orbitrap Fusion Lumos mass spectrometer (Thermo Fisher Scientific). The peptides were separated using a 210-minute gradient with a flow rate of 300 nL/min.

The mass informatics platform Proteome Discoverer 2.0 was used for post-acquisition analysis of the raw mass spectrometry data, with searches conducted using the SEQUEST HT search engine. We set the precursor mass tolerance to 10 ppm, and set the fragment ion tolerance to 0.02 Da for the Orbitrap analyzer and 0.6 Da for the ion trap analyzer. Static modifications included Tandem Mass Tags 6 (TMT6) on lysines and N-termini (+229.163 Da) and carbamidomethylation on cysteines (+57.021 Da). Methionine oxidation (+15.995 Da) was considered a dynamic modification. A UniProt *Culex quinquefasciatus* database was then used to search the data. Parent proteins were identified and quantified using the obtained peptide identification data. The abundance of the peptides was determined employing signal-to-noise (S/N) values derived from the corrected reporter ion abundances taken from the supplier’s data sheets. Subsequently, to determine the abundance of each protein, the S/N values for all its constituent peptides were log-transformed, aggregated, and summed. Analysis of variance (ANOVA) was conducted to ascertain the significant differences among the protein abundances. The FDR was then calculated to statistically validate the results, and with an FDR < 0.05 indicating a significant difference.

### RNA-seq and proteomics combined analysis

A unified analysis of the RNA-seq and proteomics data was conducted by identifying proteins and their corresponding transcripts that showed significant differential expression. The protein Log2 fold change was calculated using the protein abundance ratios, while Fragments Per Kilobase of transcript per Million mapped reads (FPKM) values were employed to determine the transcript Log2 fold change.

### Extraction of RNA and synthesis of cDNA

The wings, legs, abdomens, thoraxes, and heads were collected from female DS and DR mosquitoes at 72 h post-eclosion (PE). The same tissues from 20 mosquitoes were placed in a tube (n = 3 tubes per tissue). Total RNA was then extracted from the tissues following the guidelines of the RNAiso Plus kit (Takara, Shiga, Japan). Subsequently, as the first step of the quantitative real-time reverse transcription PCR (qRT-PCR) protocol, a Takara PrimeScriptRT Reagent Kit was employed to convert the RNA to cDNA.

### Quantitative real-time PCR (qPCR)

The prepared cDNA was diluted with an appropriate amount of RNase-free water, followed by its use as the template for qPCR, which employed a Power SYBR Green PCR Master Mix (Applied Biosystems, Foster City, CA, USA) following the supplier’s guidelines. The reactions (20 μL) consisted of the diluted cDNA, specific primer pairs (as shown in Additional file 2: Table S1), and the Power SYBR Green PCR Master Mix. The qPCR thermal program comprised: An initial denaturation at 95 °C for 10 min; and 40 cycles of denaturation at 95 °C for 15 s and annealing/extension at 60 °C for 1 min. For qPCR validation, melting curve analysis was performed immediately post-qPCR to ensure the presence of a curve with a single peak. Amplification signals in the primer control or no-template control samples exhibited high cycle threshold (Ct) values (Ct > 35). Additionally, the specificity of the primers was confirmed by sequencing the qPCR products when the primers were used for the first time. In each test, the calibration curves exhibited correlation coefficients exceeding 0.99. The internal control gene *ACTB* (encoding β-actin) was used to normalize the relative expression levels [50, 51] according to the 2−ΔΔCt method, where the formula target gene / *ACTB*= 2ΔCt was applied, with ΔCt = Ct (*ACTB*) − Ct (target gene) [52]. For qPCR analyses, three technical and biological replicates were conducted.

### Gene silencing

Female mosquitoes from the DR and DS strains were used for RNA interference (RNAi) experiments, with microinjection performed at 12 h PE (n = 3 tubes per group; 10 RNAi mosquitoes in each tube). GenePharma (Shanghai, China) synthesized small interfering RNAs (siRNAs) targeting *CYP325G4* (siCYP325G4) and *CYP6AA9* (siCYP6AA9), together with a non-targeting negative control siRNA (siNC) (Table S1). The siNC, does not induce gene silencing because it has no homologous gene targets in the mosquito genome. Approximately 364 ng of siCYP325G4, 364 ng of siCYP6AA9, and 350 ng of siNC were injected individually into female mosquito thoraxes. The detailed procedures of the gene silencing technique were carried out according to a previously published method [53]. At 3 days after injection, qRT-PCR was conducted to assess the efficiency of RNAi for the target gene.

### P450 activity

A steel pestle was employed to homogenize individual deep-frozen adult mosquitoes, guts and legs in 300 µl of cold 0.0625 M phosphate buffer (pH 7.2) at 4 °C in a flat-bottom 96-well microtiter plate. The homogenates were then subjected to centrifugation at 1109 × *g* for 20 min at 4 °C. The supernatant was retained as the enzyme source for subsequent reactions. In duplicate wells of a new microtiter plate, each reaction mixture comprised 20 µl of homogenate, 25 µl of 3% hydrogen peroxide, 200 µl of 3,3′,5,5′-tetramethylbenzidine (TMBZ) solution (0.01 g of TMBZ dissolved in a mixture of 15 ml of 0.25 M sodium acetate buffer (pH 5.0) and 5 ml of methanol), and 80 µl of 0.0625 M potassium phosphate buffer (pH 7.2). The plate was left at room temperature for 2 h and then the absorbance at 450 nm was measured as an endpoint. Equivalent units of cytochrome (EUC) P450s per milligram of protein was used to quantify the enzyme contents. A standard curve generated using purified cytochrome C was used to adjust the results for the known heme content of cytochrome C and P450s. Standard concentrations (0.01, 0.02, 0.04, 0.06, 0.08, and 0.1 mg/mL) of cytochrome C were measured and the results were plotted as a graph using Microsoft Excel (Redmond, WA, USA). Using the standard curve, the concentration of P450 monooxygenase in a sample was determined and expressed as cytochrome P450 per minute per milligram of protein. To determine the protein concentrations, in a 96-well plate, 5 µl of protein standards at various concentrations were added to separate wells, while 5 µl of the test sample was added to the sample well, followed by the addition of 250 µl of G250 staining solution to each well. After a 5 min incubation at room temperature, the absorbance of each well at 595 nm was determined. A standard curve constructed using bovine serum albumin was used to covert the absorbance values into protein concentrations. For all the biochemical assays, a minimum of three blank replicates (using water instead of the enzyme solution) were set up using the corresponding reagents and solutions for the assay. The average optical densities (ODs) of the blank replicates were subtracted from the ODs of the test wells for adjustment. The protein concentration in the sample was determined by referencing the standard curve and considering the sample volume used.

### Centers for Disease Control and Prevention (CDC) bottle bioassay

The bottle bioassay of the Centers for Disease Control and Prevention had been described earlier[47]. In each bottle, approximately 20 4-day-old non blood fed female mosquitoes from the siCYP325G4, siNC, and DEPC water groups were introduced into bottles coated with deltamethrin (0.1mg/ml) and incubated for 2 hours. Bottles coated with acetone are used as insecticide free controls. Mortality rate were assessed every 15 minutes during exposure. Each group is repeated three times.

### Scanning electron microscopy (SEM)

To account for the effect of mosquito size on cuticle thickness, the wing lengths of all female mosquitoes involved in the study were measured. Each group (siNC, siCYP325G4, and siCYP6AA9) consisted of 7–8 female mosquitoes, from which one right front leg from each mosquito was selected. The selected legs underwent two washes using 70% ethanol for thorough cleaning. Subsequently, alcohol was applied to tarsomere I of the leg at the midpoint, followed by precise cutting using a platinum-coated blade. Following cutting, the leg sections were washed to eliminate any debris. Subsequently, the sections were immersed in 2.5% glutaraldehyde (Sigma, St. Louis, MO, USA) for 12 h, followed by sequential incubation in a series of graded ethanol concentrations (30, 50, 70, 80, 90, 95, and 100%) for 10 min each time. Afterwards, an EM CPD300 critical point dryer (Leica, Wetzlar, Germany) was used to dry the sections, employing an automated process involving 15 exchanges. Finally, a K550 X sputter coater (Electron Microscopy Sciences, Hatfield, PA, USA) was utilized to coat the sections, followed by SEM analysis. To examine the cuticle thickness, SEM image analysis was conducted using Image J software (NIH, Bethesda, MD, USA). The cuticle thickness at 23 randomly selected positions was used to determine the average cuticle thickness of each leg.

### Statistical considerations

Statistical analysis was conducted using SPSS 23.0 and GraphPad Prism 6.0 software [54]. Student’s t-test was employed to compare the data between two groups. ANOVA was used to analyze the expression levels of *CYP325G4* and *CYP6AA9* in various tissues. The Chi-square test was utilized to analyze mosquito mortality[55, 56]. All results are displayed as the mean ± standard deviation (SD), and statistical significance was considered at *p* < 0.05. Each experiment was replicated in at least three independent cohorts.

## Additional files

**Additional file 1: Table S1. Primers used for qPCR analysis and siRNA synthesis of CYP325G4 and CYP6AA9.**

**Additional file 2: Summary of the transcriptome data of DR and DS legs.**

**Additional file 3: Summary of the proteome data of DR and DS legs.**

**Additional file 4: Summary of the transcriptome and proteome of DR and DS legs.**

**Additional file 5: KEGG_enrichment of the transcriptome**

**Additional file 6: KEGG_enrichment of the proteome**

**Additional file 7: KEGG_enrichment of the transcriptome and proteome**

## Abbreviations

Cx. Pipiens: Culex pipiens
iTRAQ: isobaric tags for relative and absolute quantitation
DEGs: differentially expressed genes
DEPs: differentially expressed proteins
DR: deltamethrin-resistant
DS: deltamethrin-susceptible
PE: post-eclosion
qRT-PCR: quantitative real-time reverse transcription PCR
FDR: False Discovery Rate
FPKM: Fragments Per Kilobase Million
siCYP325G4: small interfering RNA for silencing the CYP325G4 gene
siCYP6AA9: small interfering RNA for silencing the CYP6AA9 gene
NC: negative control
DEPC: diethyl pyrocarbonate
CDC: Centers for Disease Control and Prevention
SEM: scanning electron microscope
LC_50_: 50% lethal concentration
CHC: chlorinated hydrocarbon
CP: cuticle protein)
ACC: acetyl-coenzyme A carboxylase
FAS: fatty acid synthase S-acetyltransferase
FACVL: very-long-chain fatty acid-coenzyme A ligase

## Acknowledgements

Not applicable.

## Ethics approval and consent to participate

All animal procedures performed in this study were approved by the Institutional Animal Care and Use Committee (IACUC) of Nanjing Medical University. The study was conducted according to the guidelines for the use of laboratory animals, under Protocol No. 582/2017.

## Consent for publication

Not applicable

## Availability of data and materials

Data supporting the conclusions of this article are included within the article and its supplementary files. All data are fully available without restriction and can be obtained upon request.

## Authors’ contributions

YX, JJD, FFZ, XXL, LT and YFM performed experiments. YX, YC, KWZ, LM and YS wrote the manuscript and prepared the figures. FMZ, BS, YS, DZ, GYY and CLZ conceived the study and coordinated the project. All authors have read and approved the final version of the manuscript for submission.

